# Beyond the labeled line: variation in visual reference frames from intraparietal cortex to frontal eye fields and the superior colliculus

**DOI:** 10.1101/124628

**Authors:** V. C. Caruso, D. S. Pages, M. A. Sommer, J. M. Groh

## Abstract

We accurately perceive the visual scene despite moving our eyes ~3 times per second, an ability that requires incorporation of eye position and retinal information. We assessed how this neural computation unfolds across three interconnected structures: frontal eye fields (FEF), intraparietal cortex (LIP/MIP), and the superior colliculus (SC). Single unit activity was assessed in head-restrained monkeys performing visually-guided saccades from different initial fixations. As previously shown, the receptive fields of most LIP/MIP neurons shifted to novel positions on the retina for each eye position, and these locations were not clearly related to each other in either eye- or head-centered coordinates (hybrid coordinates). In contrast, the receptive fields of most SC neurons were stable in eye-centered coordinates. In FEF, visual signals were intermediate between those patterns: around 60% were eye-centered, whereas the remainder showed changes in receptive field location, boundaries, or responsiveness that rendered the response patterns hybrid or occasionally head-centered. These results suggest that FEF may act as a transitional step in an evolution of coordinates between LIP/MIP and SC. The persistence across cortical areas of hybrid representations that do not provide unequivocal location labels in a consistent reference frame has implications for how these representations must be read-out.

**New & Noteworthy:** How we perceive the world as stable using mobile retinas is poorly understood. We compared the stability of visual receptive fields across different fixation positions in three visuomotor regions. Irregular changes in receptive field position were ubiquitous in intraparietal cortex, evident but less common in the frontal eye fields, and negligible in the superior colliculus (SC), where receptive fields shifted reliably across fixations. Only the SC provides a stable labelled-line code for stimuli across saccades.

## INTRODUCTION

The perceived locations of visual stimuli are derived from a combination of the location of retinal activation and information about the direction of eye gaze. How these signals are combined to synthesize a representation of visual space as the eyes move is unknown. The process is computationally complex because any site on the retina could correspond to any given site in the visual scene, but only the correct correspondence for a particular eye gaze position is operative at any given time. Coordinate transformations are therefore both flexible and precise, and it has been suggested that they unfold as a gradual process across multiple brain regions.

Which visual areas are truly retinotopic, or eye-centered, and which employ higher-order representations incorporating information about eye movements is uncertain. The retina is thought to provide the brain with an eye-centered map of the locations of visual stimuli. But after that, the recurrent interconnectivity of the brain in principle permits adjustment of reference frame. Several studies have indicated effects of eye position or movement on visual responses as early as the lateral geniculate nucleus (Lal R and MJ Friedlander 1989, 1990, 1990). In V1, some studies have found evidence that eye position modifies visual signals (Trotter Y and S Celebrini 1999) and some have not (Motter BC and GF Poggio 1990; Gur M and DM Snodderly 1997). Later visual structures exhibit more extensive sensitivity to eye position/movements (e.g. Squatrito S and M Maioli 1996; Bremmer F et al. 1997; Bremmer F 2000).

We focus here on a quantitative comparison of the reference frames employed in three interconnected brain regions involved in guiding saccadic eye movements to visual targets (Figure 1): the lateral and medial banks of the intraparietal sulcus (LIP/MIP), the superior colliculus (SC), and the frontal eye fields (FEF) (e.g. Wurtz RH and ME Goldberg 1971; Sparks DL et al. 1976; Bruce C and M Goldberg 1985; Huerta MF et al. 1986; Lee C et al. 1988; Stanton GB et al. 1988; Waitzman DM et al. 1988; Schall J 1991; Schall J and D Hanes 1993; Freedman EG and DL Sparks 1997; Wurtz RH et al. 2001; Scudder CA et al. 2002; Bisley JW and ME Goldberg 2003; Steenrod SC et al. 2013). Electrical stimulation in the SC and FEF evokes short latency saccades at low thresholds (Robinson DA and AF Fuchs 1969; Robinson DA 1972; Schiller P, H. and M Stryker 1972; Bruce CJ et al. 1985). Stimulation in parietal cortex can also evoke saccades (Thier P and RA Andersen 1996, 1998; Constantin AG et al. 2007; Constantin AG et al. 2009). Lesions of both the SC and FEF together eliminate saccades whereas lesions of parietal cortex can produce hemineglect (Bisiach E and C Luzzatti 1978; Schiller P et al. 1980; Schiller PH et al. 1987; Sommer MA and EJ Tehovnik 1997; Dias EC and MA Segraves 1999).

Parietal cortex was one of the first brain regions in which eye movements and visual signals were shown to interact (Andersen RA et al. 1993). These response patterns were initially characterized as “gain fields”, in which receptive fields were stable in eye-centered location but exhibited a response modulation with different eye positions (Andersen RA et al. 1985). Subsequent studies involving complete sampling of receptive field positions as the eyes moved suggested that receptive fields could also adopt new positions on the retina at different eye positions (Mullette-Gillman OA et al. 2005, 2009). These changes in receptive field position produced a code that varies across neurons and ranges from predominantly eye-centered to predominantly head-centered, with most neurons exhibiting “hybrid” response patterns. In contrast, the SC, while exhibiting gain fields (Van Opstal AJ et al. 1995), employs a predominantly eye-centered code when tested and analyzed the same way as LIP/MIP (Lee J and JM Groh 2012) (see also Klier EM et al. 2001; DeSouza JF et al. 2011; Sadeh M et al. 2015).

Considerable interest has focused recently on eye position gain fields in the FEF (Cassanello CR and VP Ferrera 2007) and on how receptive fields in the FEF change transiently at the time of the eye movement (Sommer MA and RH Wurtz 2006; Zirnsak M and T Moore 2014; Zirnsak M et al. 2014). In addition, a detailed quantitative assessment of torsional, eye-in-head, and head-on-body components of the FEF’s reference frame has been conducted in a paradigm in which the both the head and eyes were free to move (Keith GP et al. 2009; Sajad A et al. 2015; Sajad A et al. 2016). However, a quantitative assessment of the reference frame during steady fixation with the head restrained (important for our larger purpose of comparing visual and auditory coding) has not, to our knowledge, been conducted.

Here we report that the reference frame of visual signals in the FEF is intermediate between the SC and LIP/MIP. The results support the view that reference frames evolve along brain pathways involved in controlling visually-guided behavior, becoming a plausible labeled line for eye-centered stimulus location only at the level of the SC.

**Figure 1:**
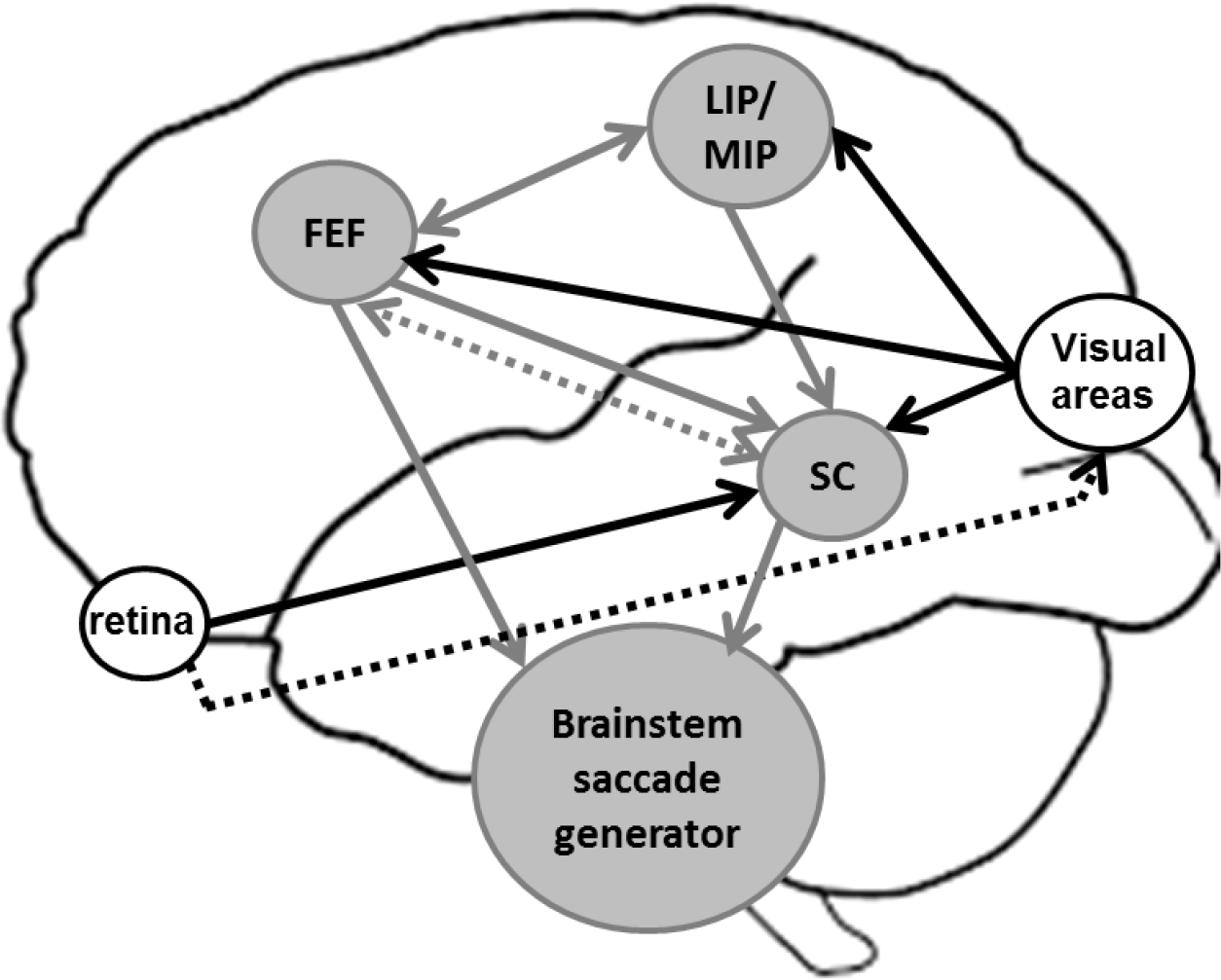
Schematic of the connections between the areas LIP/MIP, SC and FEF, their visual inputs and their projections to the brainstem saccade generator. The LIP/MIP and FEF are highly interconnected and send excitatory projections to the intermediate and deep layers of the SC (continuous arrows indicate direct projections). The FEF also sends inhibitory indirect projections to the SC through the caudate and the substanta nigra pars reticulata (dotted arrows indicate indirect projections). Both the SC and the FEF directly project to the various areas of the brainstem saccade generator system. The LIP/MIP and the FEF receive visual inputs from extrastriate visual areas. The SC receives visual inputs mainly in its superficial layer from the primary and secondary visual cortices and the FEF, and also directly from the retina. (for reviews, see Sparks DL and R Hartwich-Young 1989; Blatt GJ et al. 1990; Schall JD et al. 1995). Connections between oculomotor areas are in grey and visual inputs are in black.

## MATERIALS AND METHODS

The task and recording conditions for the FEF dataset have been explained previously (Caruso VC et al. 2016). We briefly report them here. All experimental procedures conformed to NIH guidelines (2011) and were approved by the Institutional Animal Care and Use Committee of Duke University. Two adult rhesus monkeys (*Macaca mulatta*) were implanted with a head holder to immobilize the head, a scleral search coil to track eye movements and a recording cylinder over the left or right FEF. Similar procedures were used to prepare for recordings in LIP/MIP and SC, as reported previously (Mullette-Gillman OA *et al*. 2005, 2009; Lee J and JM Groh 2012, 2014)

All data were recorded while the monkeys performed visually or aurally guided saccades, randomly interleaved, in an overlap saccade paradigm. Only visual trials were analyzed in this study. In each trial, a target was presented while the monkey fixated a visual fixation stimulus (Figure 2A, B; all visual stimuli were produced by green light emitting diodes, LEDs). The monkey withheld the saccade for 600-900 ms until the offset of the fixation target, permitting the dissociation of sensory-related activity from motor-related activity. The targets were located in front of the monkeys at 0° elevation and between -24° and *24° horizontally (nine locations separated by 6°, figure 2A). In each session, all saccades started from three initial fixation locations at -12°, 0°, *12° along the horizontal direction and at an elevation chosen to best sample the receptive field of the neuron under study.

The behavioral paradigm, the acquisition of eye trajectory and the recordings of single cell activity were controlled using the Beethoven program (Ryklin Software). Eye gaze was sampled at 500Hz. Single neuron extracellular activity was acquired using a Plexon system (Sort Client software, Plexon) through tungsten micro-electrodes (FHC, 0.7 to 2.5 MOhm at 1 kHz).

### Analysis

All analyses were conducted with custom-made routines in Matlab (the MathWorks Inc.). Only correct trials were included.

*Spatial selectivity analysis:* This analysis has been described in detail in (Caruso VC *et al*. 2016), (Mullette-Gillman OA *et al*. 2005) and (Lee J and JM Groh 2012). Briefly, we defined a **baseline period**, comprising the 0-500 ms of fixation before the target onset and a **sensory window** as the period 0-500 ms after target onset. The sensory window captured both the transient and sustained responses to visual targets (We selected the **motor window** differently in different areas, to better capture the saccade related burst, which has different temporal characteristics across regions. The motor window was defined to start before the saccade onset (20ms before saccade onset for the SC data, 50ms before for the FEF data and 150ms before for the LIP data) and to end at saccade offset. Saccade onset and offset were defined as the moment, at 2 ms resolution, that the instantaneous speed of the eye movement exceeded or dropped below a threshold of 25°/s. (In addition to these fixed analysis windows, we also analyzed sliding 100 ms windows throughout the interval from target onset to the saccade, detailed below at the end of this section).

Neurons were considered **responsive** in the sensory/motor intervals if a two-tailed t-test between their baseline activity and relevant response period was significantly different with p<0.05. Spatial selectivity of responses (in the sensory or motor period) was assessed in both head- and eye-centered reference frames, using two two-way ANOVAS. Each ANOVA involved the three levels of initial eye position (-12°, 0°, *12°) as well as five levels of target location (-12° to *12° in 6° increments), defined in head-centered coordinates for the first ANOVA and in eye-centered coordinates for the second ANOVA. Cells were classified as **spatially selective** if either of the two ANOVAs yielded a significant main effect for target location, or a significant interaction between the target and fixation locations (Mullette-Gillman OA *et al*. 2005, 2009; Lee J and JM Groh 2012, 2014). In all tests, statistical significance was defined as p value < 0.05. To be consistent with our previous analyses and because these tests were used for inclusion criteria rather than hypothesis testing, we did not apply Bonferroni correction.

*Reference frame analysis:* To distinguish eye-centered and head-centered reference frames, we quantified the degree of alignment between eye-centered and head-centered tuning curves obtained from trials with initial eye positions at -12º, 0 º, *12º along the horizontal axis. This analysis was applied to single cells during different time windows throughout the trials. In particular, for each time window considered, we constructed the three response tuning curves for the three fixation locations with target locations defined in head- or eye-centered coordinates (schematized in figure 2) and quantified their relative shift with an index akin to an average correlation coefficient (equation 1, (Mullette-Gillman OA *et al*. 2005)). We call it **reference frame index** and for each response we calculate two indexes, in head-centered and in eye-centered coordinates, according to the formula:

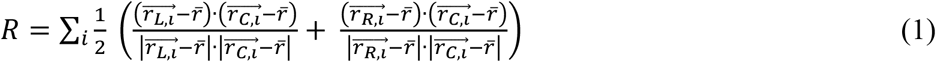

where r_L,i_, r_C,i_, and r_R,i_ are the vectors of average responses of the neuron to a target at location *i* when the monkey’s eyes were fixated at the left (L), right (R), or center (C). Only the target locations that were present for all three fixation positions in both head- and eye-centered frames of reference were included (5 locations: -12, -6, 0, 6, and 12°). The reference frame index is primarily sensitive to the relative translation of the three tuning curves and is comparatively insensitive to possible gain differences between them, provided the sampling includes some inflection point in the response curve. This can occur either by sampling from both sides of the receptive field center or by sampling locations that are both inside and outside of the receptive field. The reference frame index values range from -1 to 1: 1 indicates perfect alignment, 0 indicates no alignment, -1 indicates perfect negative correlation (Mullette-Gillman OA *et al*. 2005; Porter KK and JM Groh 2006; Mullette-Gillman OA *et al*. 2009; Lee J and JM Groh 2012, 2014). We calculated the 95% confidence intervals of the index with a bootstrap analysis (1000 iterations of 80% of data for each target/fixation combination). Each set of responses was classified based on the quantitative comparison between the eye- and head-centered indices as: 1) **eye-centered**, if the eye-centered reference index was statistically higher than the head-centered reference index (that is: if the 95% confidence interval of eye-centered reference index was positive and larger than the 95% confidence interval of head-centered reference index); 2) **head-centered**, if the opposite pattern was found; 3) **hybrid-partial shift**, if the two indexes were statistically different from each other, and at least one of them was statistically different from zero (that is: if the 95% confidence intervals of the eye-and head-centered indices overlapped with each other and at least one did not include zero) **4) hybrid-complex**, if the two indexes were not statistically different from each other and from zero (that is: if the 95% confidence intervals of the eye-and head-centered indices overlapped with each other and with zero) (Figure 4). These latter two categories were combined in our previous reference frame analyses of activity patterns in the SC and LIP/MIP. (Mullette-Gillman OA *et al*. 2005, 2009; Lee J and JM Groh 2012). Here, we consider them both separately and together as appropriate to provide a more comprehensive comparison of the reference frames across regions. We conducted the reference frame analysis for each cell during the sensory and motor periods (figure 4A,B and 5A,B) and across time in 100ms windows sliding with steps of 50ms from the target onset and from saccade onset (figure 4C,D, and 5C,D).

### LIP/MIP and SC datasets

Figure 5 includes data from area LIP/MIP and SC that we have previously collected and described (Mullette-Gillman OA *et al*. 2005, 2009; Lee J and JM Groh 2012, 2014). The tasks and recording techniques were the same as in the present study, but each dataset was recorded from different monkeys (two monkeys per brain area). The SC dataset consists of a total of 179 single cells recorded in the left and right SC of two monkeys. The LIP/MIP dataset consists of a total of 275 single cells recorded in the left and right LIP/MIP of two monkeys. For these two additional datasets, responsiveness, spatial selectivity and reference frame were assessed the same way as for the FEF data set. Some of the data in figure 5 were reanalyzed in different time windows than the original studies, to allow for a fair comparison across brain areas. However, changing the analyses windows did not change the overall results of the previous studies.

## RESULTS

### Overview

We first describe our new data concerning the visual reference frame in FEF before quantitatively comparing FEF to our previously reported results in LIP/MIP and the SC ((Mullette-Gillman OA *et al*. 2005, 2009; Lee J and JM Groh 2012, 2014)). We recorded single cell response profiles while the monkeys performed visually guided saccades, interleaved with aurally guided saccades which were not analyzed here (see Methods). The task and the location of the stimuli are described in figure 2A,B.

We tested whether the receptive fields of FEF neurons (N=324) shifted with the eyes or stayed fixed relative to the head by defining two reference frame index R_eye_ and R_head_ in eye- and head-centered coordinates. These indexes are akin to the average correlation coefficient between the tuning curves obtained from the three initial fixations (Methods). Thus they measure the relative alignment of the tuning curves in eye-centered or head-centered coordinates, without regard to changes in overall response magnitude, which does not contribute substantially to this measure. Figure 2 schematically shows our classification system based on the quantitative comparison between R_eye_ and R_head_. Response patterns were considered either **“eye-centered”** (Figure 2C: the tuning curves aligned in eye-coordinates better than in head-coordinates), **“head-centered”** (Figure 2D: opposite pattern), or **“hybrid”** if the response patterns were not well described in either coordinate system. A hybrid reference frame refers to circumstances such as when the retinal positions of receptive fields differ across eye positions but not in the way consistent with head-centered coordinates, either because the receptive fields shift by only part of the difference in fixation position (Figure 2E, we refer to this scenario as **“hybrid-partial shift”**) or because the receptive fields change across fixations in unpredictable ways that appear random (Figure 2F, **“hybrid-complex”**). Figure 2 G-J shows the patterns expected when R_head_ is plotted against R_eye_ in each of these scenarios: data points would cluster below (G), above (H) or along the diagonal (I,J) depending on the observed coordinate system.

A final note is that in these experiments, head, body, and world are immobile with respect to each other. Thus it is not possible to distinguish these reference frames from one another.

**Figure 2:**
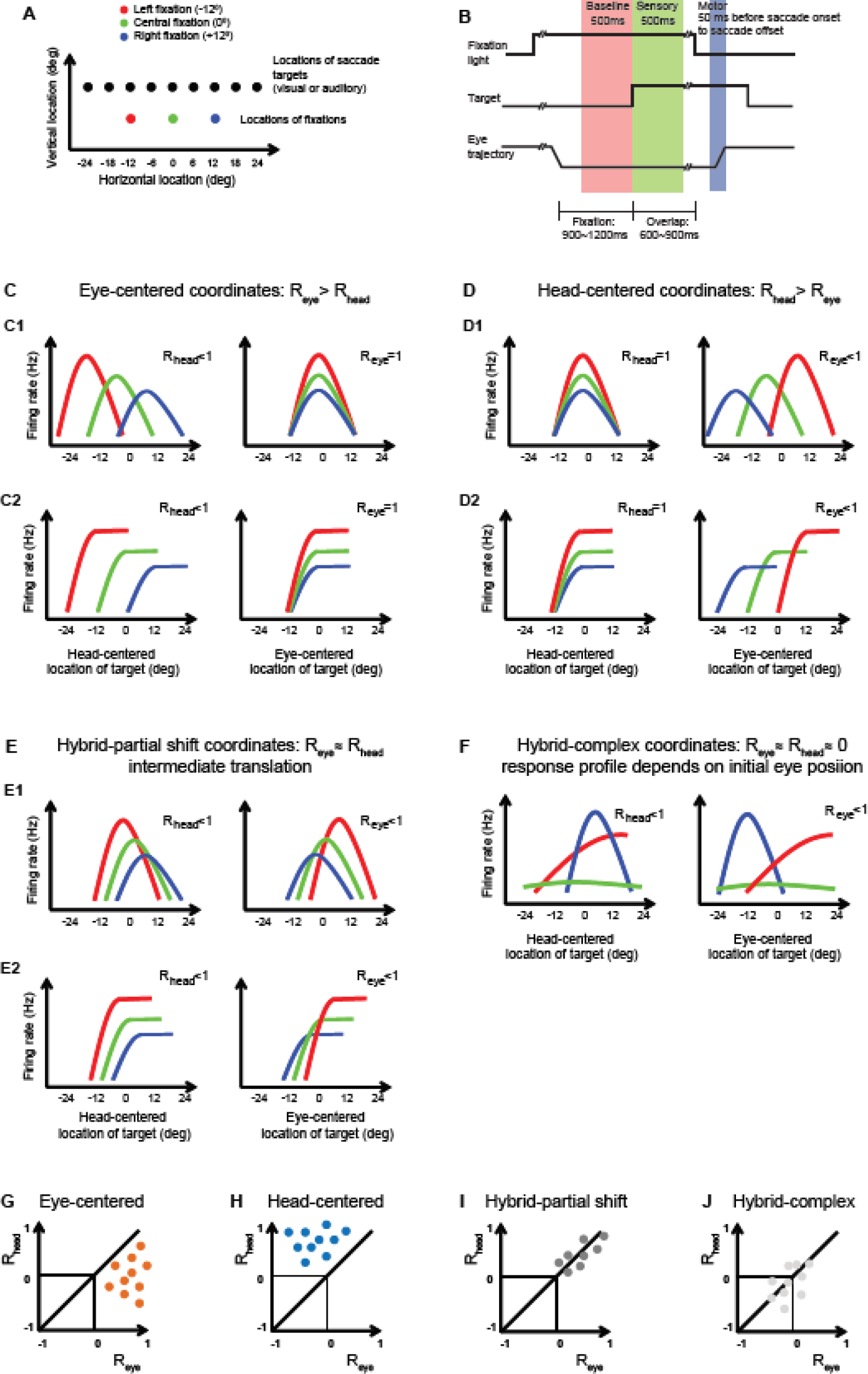
Stimuli, task and classification system of receptive fields (A) Locations of stimuli and initial fixations: Varying the initial fixation permits the separation of eye- and head-centered reference frames by measuring the relative alignment of the responses in head- and eye-centered coordinates. **(B) Task**: Each trial starts with the appearance of a fixation light, which the monkey is required to fixate. A target then appears, but the monkey needs to wait until the fixation goes out bfore making a saccade to the target. **(C) Predominantly eye-centered response patterns**. The three tuning curves obtained for the three initial fixation locations align best in eye centered coordinates (perfect alignment, R_eye_≈1, right panel), while in head-centered coordinates (left panel), they are shifted by the distance between the initial eye positions (i.e. steps of 12º in the present task, resulting in R_head_<1). **(D) Predominantly head centered responses**. The pattern is the opposite of (C): the three tuning curves are aligned in head-centered coordinates and separated by 12º in eye-centered coordinates. **(E) Hybrid-partial shift response pattern**. The three tuning curves are not well aligned in either head- or eye-centered coordinates: as the initial eye direction shifts left or right (in red and blue), the tuning curves only partially move apart, by less than 12º. **(F) Hybrid-complex coordinates**. The initial eye location affects the shape, the gain and/or the alignment of the tuning curves in unpredictable ways that have no obvious relationship in either eye- or head-centered coordinates. **(G-J) Schematics of population analysis.** When R_head_ is plotted vs. R_eye_, the data points should lie below the line of slope one if the reference frame is predominantly eye-centered **(G)**, above the line of slope one if head-centered **(H)**, along the line of slope one, but at positive values, if hybrid-partial shift **(I)** and should look random if hybrid-complex **(J)**.

### Example Neurons in the FEF

Figure 3 shows the responses from nine individual cells during the sensory (Figure 3 A-C-E-G-I) and motor periods (Figure 3 B-D-F-H-J). During both the sensory period and the motor burst, the most common pattern was eye-centered (Figure 3A-B-C-D) with an unambiguous difference between high reference frame indexes in eye-centered coordinates (close to 1), and very low reference frame indexes in head-centered coordinates (the examples shown in Figure 3A-B-C-D had, respectively: R_eye_ = 0.97; 0.93; 0.82 and 0.97 vs. R_head_ = -0.17; 0.18; 0.37 and 0.35).

Some neurons were classified as head-centered on the basis of the quantitative comparison of the reference frame indexes (R_head_>R_eye_), but as can be seen in Figure 3E-F, these response patterns were not as strongly head-centered as the eye-centered neurons were eye-centered. The values of R_head_ for these two examples were 0.68 and 0.58, whereas the values of R_eye_ were 0.31 and 0.28.

There were also neurons that exhibited hybrid response patterns. The example in Figure 3G shows hybrid-complex tuning curves: two of the curves are well aligned in head-centered coordinates (the red and green ones, representing the receptive fields computed from the left (red) and central (green) fixation location), and two are well aligned in eye-centered coordinates (the green and blue ones, representing the receptive fields computed from the center and right fixation location). The example in Figure 3H shows hybrid-partial shift tuning curves: the curves clearly shift laterally when the eyes move to different initial fixation location, but not by the correct amount. In both of these examples, R_eye_ and R_head_ are about equal to each other (Figure 3G: R_eye_=R_head_=0.34; Figure 3H: R_head_=0.81, R_eye_=0.80).

A final response pattern involved responses that were only weakly modulated by target location in any reference frame. Figure 3I-J illustrates examples in which responses exceeded baseline for all locations tested, but there was little evidence of a circumscribe dreceptive field among the locations tested. Target-evoked responses even occurred for locations well into the ipsilateral field of space. Note that these responses were not necessarily identical for all locations, and could vary with eye position (e.g. the curves in Figure 3I-J show an overall gain sensitivity to eye position). However, the lack of spatial sensitivity in these neurons makes it impossible to evaluate their reference frame. Accordingly, we tested for spatial sensitivity using an ANOVA involving the 5 targets at-12º, -6 º, 0 º, 6 º, 12º (see Methods and Results-Overview). Neurons that failed to show spatial sensitivity in either head- or eye-centered coordinates made up about half of the sample in FEF (as well as LIP/MIP), and these neurons were excluded from the population analyses described in the next section.

**Table 1:** Spatially selective populations in LIP/MIP, FEF, SC.

|  | Number of<br>recorded cells | Spatially<br>selective cells<br>in the sensory<br>period | Spatially<br>selective cells<br>in the motor<br>period | Spatially<br>selective cells<br>in the sensory<br>and motor<br>period |
| --- | --- | --- | --- | --- |
| LIP/MIP | 275 | 125 (45%) | 127 (46%) | 69 (25%) |
| FEF | 324 | 174 (54%) | 160 (49%) | 117 (36%) |
| SC | 179 | 162 (91%) | 161 (90%) | 153 (85%) |

*Table 1 legend: For each area (LIP/MIP, FEF, SC) and each time window (sensory and motor periods, see Methods) the number of spatially selectivity cells is indicated. A cell was considered spatially selective if its response was modulated by target location in either head- or eye-centered coordinates, according to two two-way ANOVAs with target location and fixation position as the two factors (one two-way ANOVA for target locations defined with respect to the eye and one two-way ANOVA for target location defined with respect to the head; target location main effects p<0.05; interaction terms p<0.05)*.

**Figure 3:**
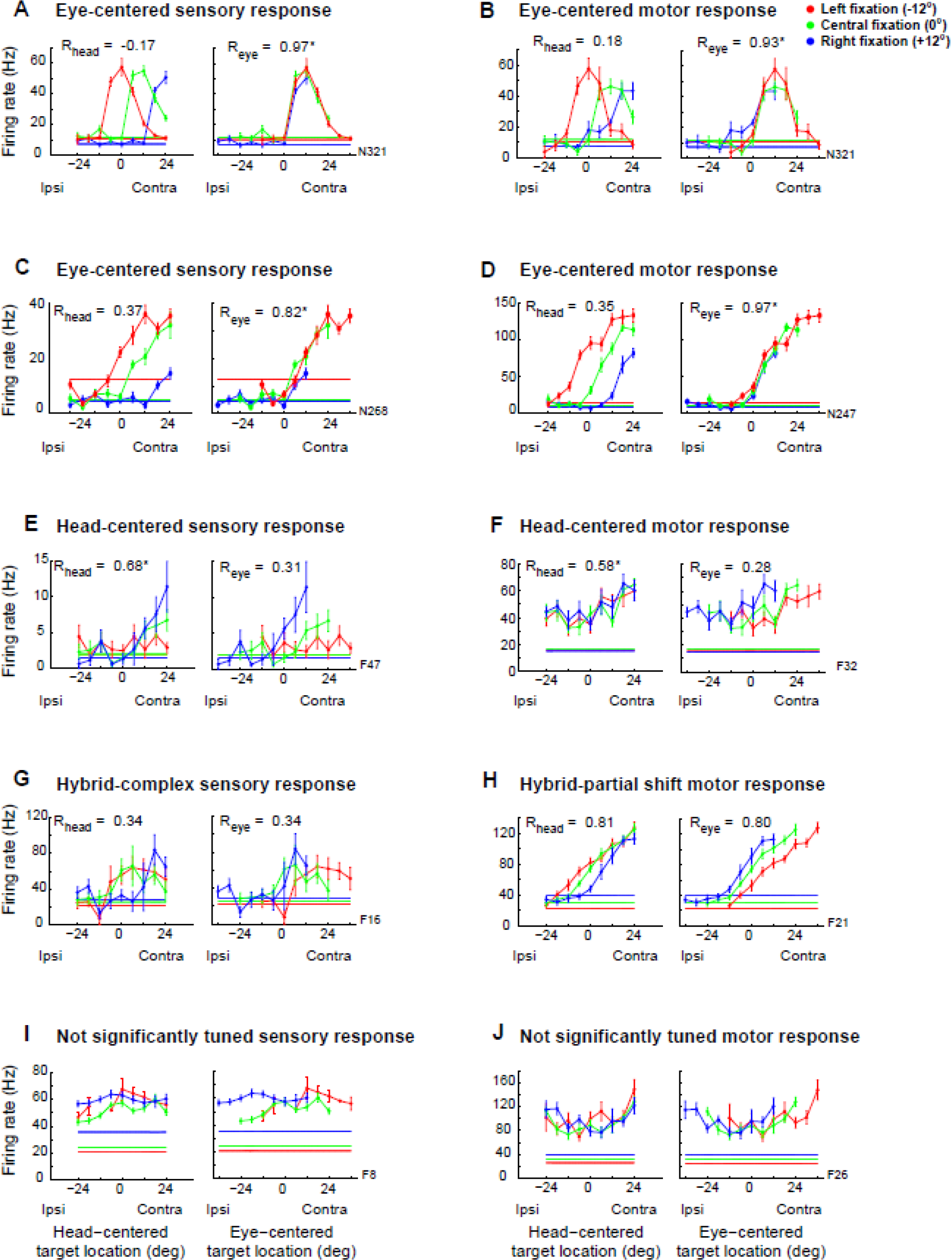
Examples of responses in the FEF. Each panel shows the tuning curves for various example cells during the sensory or motor period (Methods). The tuning curves are plotted both in head-centered coordinates (left) and eye-centered coordinates (right) and the reference frame indexes R_head_ and R_eye_ are indicated. **(A-B-C-D)** Examples of eye-centered responses (R_eye_ statistically higher than the R_head_) during the sensory period (A and C) and during the motor burst (B and D). The two responses in A and B (from the same cell at different time windows) show a complete sampling of the receptive field, while the examples in C and D show a partial sampling of the receptive fields. **(E)** Head-centered responses (R_eye_ statistically smaller than the R_head_) during the sensory period. **(F)** Head-centered responses during the motor burst. **(G)** Hybrid-complex responses (R_eye_ not statistically different from R_head_ and both not statistically different from zero) during the sensory period. **(H)** Hybrid-partial shift responses (R_eye_ not statistically different from R_head_, but at least one of them statistically higher than zero) during the motor burst. **(I)** Untuned responses during the sensory period. **(J)** Untuned responses during the motor burst. The reference frame index R was not calculated for the responses not significantly modulated by the target location as showed in panels G and H.

### Population Results in the Frontal Eye Fields

Eye-centered responses in the FEF are prevalent in both sensory and motor periods, corresponding to about 60% of the population. Figure 4A and B show this quantitatively. Like Figure 2D-F-H-J, these graphs plot R_eye_ vs. R_head_. The cross hairs indicate the 95% confidence intervals on those values (see Methods). Data points are color coded orange if the 95% confidence intervals suggest R_eye_>R_head_ (the 95% range of R_head_ is lower than the 95% range of R_eye_), blue if R_head_>R_eye_, and gray if R_eye_ and R_head_ are approximately equal; the two shades of gray correspond to R_eye_~=R_head_ and either R_eye_>0 or R_head_>0 (dark grey) and R_eye_~=R_head_ ~=0 (light grey). The pie charts on the right indicate the percentages of cells classified as eye-centered (orange), head-centered (blue), or the various subtypes of hybrid coding.

The pattern of reference frames was similar in the sensory vs. motor periods, and there was little evidence of a systematic change in the representation when time scale was investigated more closely. Figure 4C-D shows the average R_eye_ and R_head_ values in 100 ms time bins sliding in 50 ms increments from target onset (figure 4C) and from saccade onset (figure 4D). R_eye_ averages about 0.4 whereas R_head_ averages about 0.1, with little change except for slight upticks at target onset and saccade onset. Although R_eye_ is consistently significantly greater than R_head_ (filled symbols, t-test on each time bin, p<0.05), its value is not very high in absolute terms. This is because only about 60% of FEF neurons are classifiable as having eye-centered responses and among these, many R_eye_ values were not very high. On the whole, the reference frame in the FEF is more eye-centered than it is head-centered, but it is not fully eye-centered.

**Figure 4:**
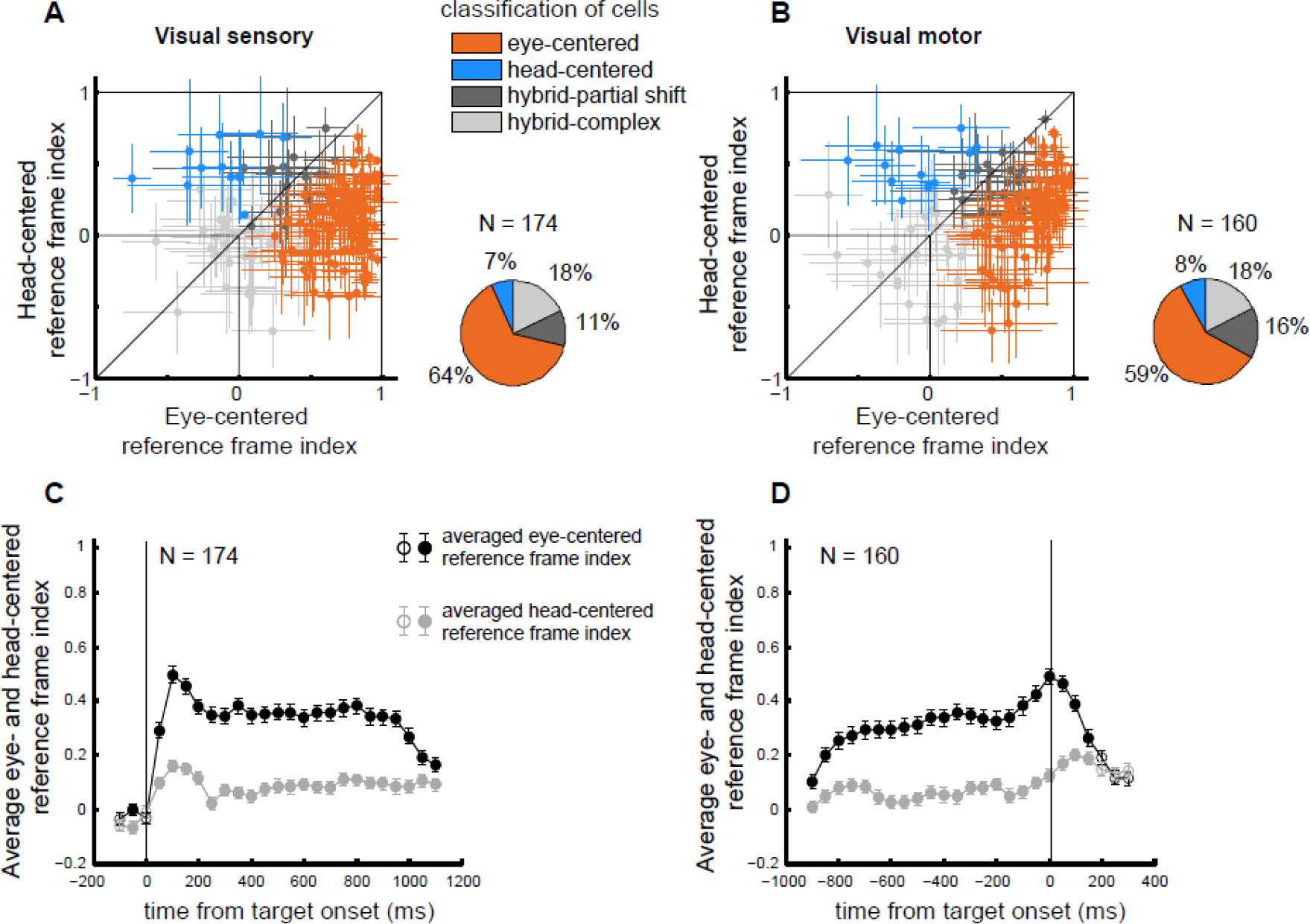
Reference frames in the FEF population. (A,B) The reference frame indexes in head-centered and eye-centered coordinates are plotted for spatially selective cells in each time window: **(A)** visual sensory, **(B)** visual motor. Responses are classified as eye centered if the 95% confidence interval of eye-centered coefficient was positive, larger than, and non-overlapping with the 95% confidence interval of head-centered coefficient (bootstrap analysis, see Methods). These are indicated in orange. Responses were classified as head centered with the opposite pattern (blue). Finally, hybrid-partial shift reference frames, in dark grey, have non-zero overlapping 95% confidence intervals, while hybrid-complex responses, in light grey, have reference frame indexes not statistically different from zero. The pie charts summarize the proportion responses classified as eye-centered, head-centered, and the subtypes of hybrid for each time window. **(C-D)** Time course of the average eye-centered (black) and head-centered (grey) reference frame indices for the FEF population of spatially selective responses (Methods). The indices were calculated in bins of 100ms, sliding with a step of 50ms and averaged across the population. Trials were aligned at target onset (**C**) and at saccade onset (**D**). Filled circles indicate that the difference between the two average indexes is statistically significant (t-test for each bin, p<0.05).

### Comparison with LIP/MIP and the SC

We next asked how the degree of eye-centeredness in the representation in the FEF compares with the representations in parietal cortex and the SC. Figure 5 presents a comparison of the signals observed in each brain region under identical experimental conditions (see Methods and (Mullette-Gillman OA *et al*. 2005, 2009; Lee J and JM Groh 2012, 2014) for the description of the two additional datasets recorded in different monkey with the same technique and during the same task). As in the analysis of the FEF, we focus on two measures. First, we compare the number of cells classified as eye-centered, head-centered, or hybrid across areas (figure 5A, B). Second, we compared the time course of the population-averaged eye-centered reference frame in the LIP/MIP, FEF, SC (figure 5C, D).

The results indicate a continuum of reference frame across these brain areas, with LIP/MIP predominantly hybrid, the SC predominantly eye-centered, and FEF intermediate between the two. In both sensory and motor periods, the proportion of eye centered responses remains a minority in the LIP/MIP (increasing from ~20% to ~40% in time) and a majority in the SC (~80% consistently in the two time periods). The FEF is between the two trends, with ~60% eye-centered responses in both time periods. Conversely, hybrid responses dominate in LIP/MIP (from ~60% to ~40%), fall to less than 40% in the FEF and to less than 20% in the SC. Head centered responses are a minority in all three areas, though a trend can be seen in both periods with the LIP/MIP having around one fifth of responses head-centered, to the FEF having fewer than 10%, to the SC having almost no head-centered signals (Figure 5A-B).

A similar picture emerges when considering the average values of the reference frame indices across time. In Figure 5CD, the average value of R_eye_ for each brain area is plotted across time (colors indicate the areas, filled dots indicate bins where the eye-centered reference frame was statistically larger than the corresponding head-centered reference frame index). Since the proportion of spatially selective cells varied in the three areas, we repeated the analysis twice: for the subpopulation of spatially selective cells (figure 5C) and for all recorded cells (figure 5D). In either population, the visual responses are significantly more eye-centered than head-centered in all areas, but with a clear grading from SC being the most eye-centered (average index around 0.6/0.8 during the trial) to FEF (average index around 0.2/0.4 during the trial, occasionally reaching 0.5) to LIP/MIP (average index consistently lower than 0.2).

**Figure 5:**
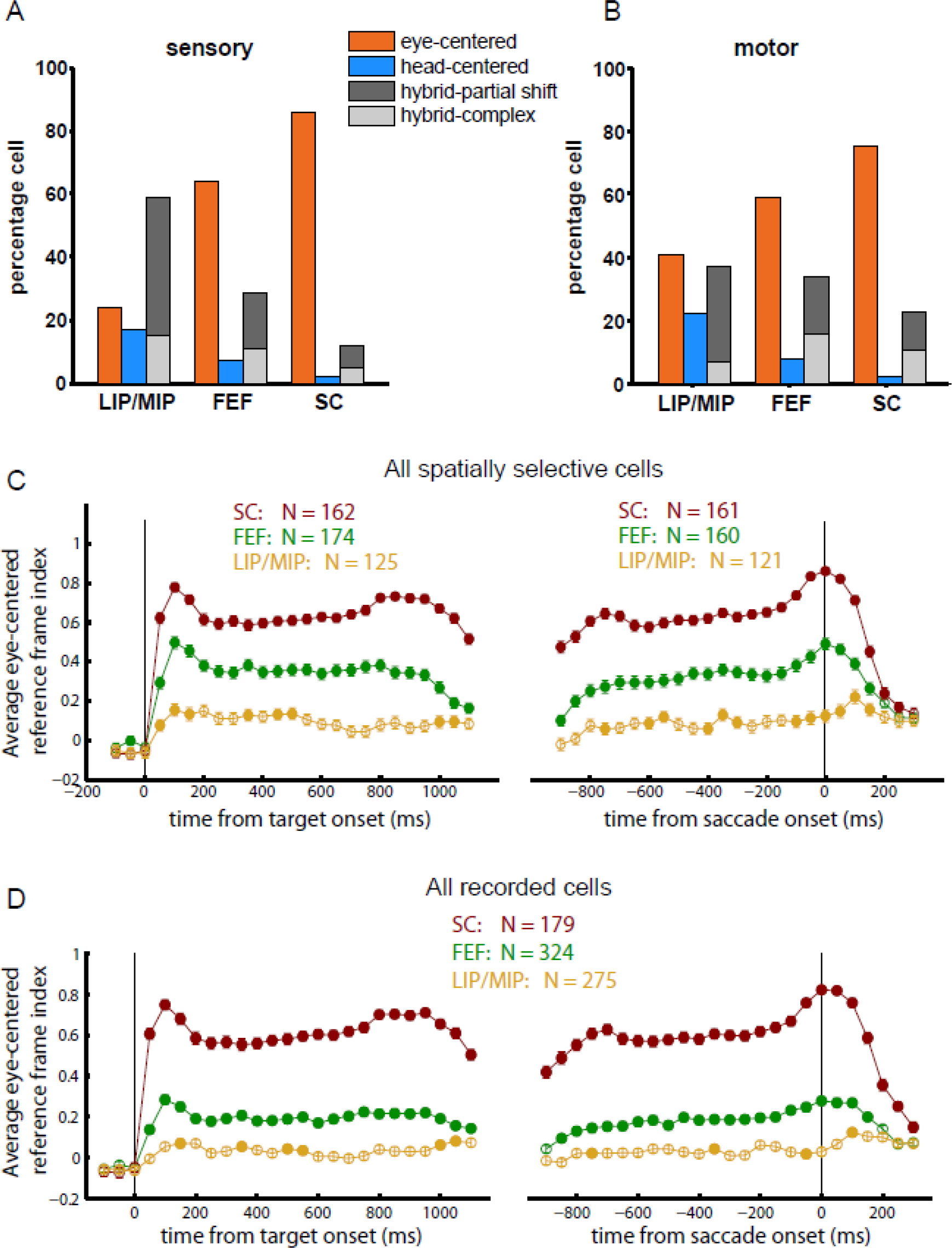
Reference frame indexes during sensory and motor periods in LIP/MIP, FEF and SC. (A-B) The percentage of cells classified as eye-centered (orange), head-centered (blue) and hybrid (gray shades: dark gray = hybrid-partial shift, light grey = hybrid-complex) are shown for all spatially selective cells in LIP/MIP, FEF and SC in the sensory period **(A)** and in the motor period **(B)**. For the sensory response, the time window is the 500 ms immediately after target onset. For the motor response, the time window changes with the saccade duration, ranging from -20ms (SC), -50ms (FEF) or -100ms (LIP/MIP) from saccade onset to saccade offset (see Methods). For the FEF, the data are the same as in figure 5A, B. **(C-D)** time course of the eye-centered reference frame indexes for the populations of spatially selective cells in LIP/MIP, FEF, SC **(C)** and for all recorded cells **(D)**. Colors: LIP/MIP (tan), FEF (green) and SC (dark red). Filled circles indicate that the eye-centered correlation coefficient was significantly larger than the head-centered one (two-tailed t-test for each bin, p<0.05). The FEF data are the same as in figure 4C. The LIP/MIP and SC data were collected in (Mullette-Gillman OA et al. 2005, 2009; Lee J and JM Groh 2012, 2014)).

## DISCUSSION

Our quantitative assessment of the reference frame for visual space representation in the FEF shows that visual signals were mostly but not completely eye-centered. Around 60% of individual neurons were classified as eye-centered, whereas one third of neurons had hybrid reference frames and a minority had weakly head-centered reference frames. At the population level, the eye-centered reference frame index averaged about 0.2-0.4, and was largely stable across time. In comparison to the SC and LIP/MIP, the pattern of reference frame in the FEF was less eye-centered than the SC but more eye-centered than LIP/MIP, which is predominantly hybrid in its coding format.

It is a challenge to compare our results to most previous studies because misinformation about the reference frames employed in the FEF and LIP/MIP pervades the literature. It is typical to find statements in abstracts or introduction sections asserting that the reference frames employed in these areas are known to be retinotopic or eye-centered, but the citations offered do not necessarily support those claims. The most apt comparisons with our experiments would be studies that (a) systematically varied initial eye position, (b) sampled receptive fields along the same dimension as the variation in eye position, and (c) quantified the results at the population level are relevant to this question. For a more detailed discussion of how the assessment of reference frame can go wrong when these conditions are not met, see Figures 10-11 of Mullette-Gillman et al (Mullette-Gillman OA *et al*. 2009).

With these caveats in mind, our FEF results are actually on the whole consistent with the most relevant previous studies. The closest related studies involved visual memory and 3D movements in head-free monkeys (Keith GP *et al*. 2009; Sajad A *et al*. 2015; Sajad A *et al*. 2016). These studies allowed natural variation in initial eye and head position and used different analysis methods to assess a variety of different reference frames. The eye-centered reference frame was the most common best match, although most other reference frames could not be statistically ruled out. Several studies involving receptive field mapping at the time of eye movements have confirmed the proof of principle that many FEF receptive fields do not maintain a fixed position on the retina across eye movements (Umeno MM and ME Goldberg 1997, 2001; Sommer MA and RH Wurtz 2006; Zirnsak M *et al*. 2014).

Studies employing electrical stimulation have also shown mixed reference frame information. Although stimulation doesn’t typically drive the eyes to a fixed position in the orbits, the direction and amplitude of the stimulation-evoked saccade can vary with initial eye position (Robinson DA and AF Fuchs 1969; Bruce CJ *et al*. 1985; Dassonville P et al. 1992; Russo GS and CJ Bruce 1993; Monteon JA et al. 2013). Monteon et al (2013) report that about 71% of sites yield saccades of a stable vector across different eye positions (i.e. eye-centered) and about 29 % yield saccades that vary considerably, more consistent with a head- or body-centered coordinate system.

The range of reference frames observed in the FEF provides a context for interpreting those observed in both the SC and LIP/MIP. The SC has been characterized as an eye-centered structure (Schiller P, H. and M Stryker 1972; Klier EM *et al*. 2001; Lee J and JM Groh 2012), while the LIP/MIP is not. Mullette-Gillman et al. (Mullette-Gillman OA *et al*. 2005, 2009) were the first to describe the reference frame of LIP/MIP as predominantly hybrid. Andersen et al. (e.g. Andersen RA and VB Mountcastle 1983; Andersen RA *et al*. 1985; Andersen RA et al. 1990) had already reported an interaction between visual inputs and eye position in the LIP/MIP, which they interpreted as reflecting eye centered gain fields. However, their sampling of receptive fields sometimes focused on the dimension orthogonal to the change in fixation position, which made their results difficult to interpret. Adopting a more appropriate sampling technique across brain areas, we have demonstrated not only strong eye-centered signals (the vast majority in the SC) but also signals along a continuum between eye-centered and head-centered coordinates (in different proportions, in all three areas). Thus, the hybrid reference frames that we have characterized are unlikely to be due to methodological shortfalls. Furthermore, the identification, under the same conditions, of a gradual shift in the strength of eye-centered representations from LIP/MIP to FEF to SC supports the view that there is a genuine transition in coding between these different brain areas. How this observation may interact with potential differences in the breadth of receptive fields across these structures remains to be determined.

Why the brain uses reference frames that are impure, and why this should vary across brain areas is unclear. The receptive field as a labelled line for stimulus location is a concept that dates back to the discovery of receptive fields. However, if receptive fields can vary in their position, and do not show stability with respect to either eye- or head-centered coordinates, such neurons on their own cannot provide a labelled line for stimulus location. Rather, the activity of a population of such neurons would be required to disambiguate the spatial signals. While this is very possible (e.g. Pouget A and TJ Sejnowski 1997), why it is desirable from a coding perspective has not been shown. Models of coordinate transformations show that the brain should be capable of transforming reference frames in a single step without the use of intermediate stages ((Groh JM and DL Sparks 1992)).

An understanding of how coordinate transformations unfold in a sensorimotor context has implications more broadly, as this is any example of a many-to-many mapping that occurs in numerous other contexts. For example, numerous types of perceptual constancy involve physical stimuli that vary but produce similar percepts (space constancy, size constancy, color constancy, the same musical melody in different keys, etc.). Such constancy can require a mapping from nearly any possible range within a physical stimulus dimension to a common perceptual dimension. Similarly, categorization requires mapping different types of physical stimuli that can be grouped into a category, and associative learning involves relating two different physical stimuli in a potentially arbitrary fashion. The underlying computation supporting these abilities may be very similar to that involved with spatial coordinate transformations.

## ACKNOWLEDGEMENT

This work was supported by the National Institute of Health R01 NS50942-05. The authors thank Jessi Cruger, Karen Waterstradt, Christie Holmes and Stephanie Schlebusch for animal care and Tom Heil for technical assistance. We have benefitted from thoughtful discussions with JungAh Lee, Kurtis Gruters, Shawn M. Willett, Jeffrey M. Mohl, David Murphy, Bryce Gessell and James Wahlberg.

